# Developmentally programmed switches in DNA replication: gene amplification and genome-wide endoreplication in Tetrahymena

**DOI:** 10.1101/2022.12.15.520636

**Authors:** Xiangzhou Meng, Hung Quang Dang, Geoffrey M. Kapler

**Affiliations:** Department of Cell Biology and Genetics, Texas A&M University Health Science Center; State Key Laboratory of Microbial Metabolism, and School of Life Sciences and Biotechnology, Shanghai Jiao Tong University, Shanghai, China; Alstem Bioscience, Richmond, CA

**Keywords:** DNA replication, endoreplication, gene amplification, macronuclear development, Tetrahymena

## Abstract

Locus-specific gene amplification and genome-wide endoreplication generate the elevated copy number of ribosomal DNA (rDNA, 9000 C) and non-rDNA (45 C) chromosomes in the developing macronucleus of *Tetrahymena thermophila*. Subsequently, all macronuclear chromosomes replicate once per cell cycle during vegetative growth. Here we describe an unanticipated, programmed switch in the regulation of replication initiation in the rDNA minichomosome. Early in development the 21 kb rDNA minichromosome is preferentially amplified from 2 C to ~800 C from well-defined origins, concurrent with genome-wide endoreplication (2 C to 8-16 C) in starved mating Tetrahymena (endoreplication (ER) Phase 1). Upon refeeding, rDNA and non-rDNA chromosomes achieve their final copy number through resumption of just the endoreplication program (ER Phase 2). Unconventional rDNA replication intermediates are generated primarily during ER phase 2, consistent with delocalized replication initiation and possible formation of persistent RNA-DNA hybrids. Origin usage and replication fork elongation are affected in non-rDNA chromosomes as well. Despite the developmentally programmed 10-fold reduction in the ubiquitous eukaryotic initiator, the Origin Recognition Complex (ORC), active initiation sites are more closely spaced in ER phases 1 and 2 compared to vegetative growing cells. We propose that initiation site selection is relaxed in endoreplicating macronuclear chromosomes and may be less dependent on ORC.

## INTRODUCTION

The conventional cell cycle of eukaryotes consists of four phases: G1, S, G2 and M. Gap phases (G1, G2) constitute the major periods for protein biosynthesis and cell growth, while S and M are devoted to DNA synthesis and chromosome segregation (mitosis or meiosis), respectively. Cell cycle regulation is imparted in large part by the biogenesis, post-translational modification and/or degradation of cyclins, cyclin-dependent kinases and proteins that are required to initiate DNA replication and segregate chromosomes (reviewed in Morgan, 1997 and in Dolson et al., 2021). Cell cycle checkpoints maintain genic balance by assuring that chromosomes are duplicated once and only once during S phase, and are properly segregated during mitosis.

Deviations from the conventional cell cycle are critical for metazoan development and increase gene dosage in a global or locus-specific manner. Genome-wide endoreplication consists of repeated Gap-S-Gap-S phases without cell division, and typically occurs in specialized terminally-differentiated cells (reviewed in Orr-Weaver, 2015 and in Shu, Row and Deng, 2018). The net result is the biogenesis of polyploid nuclei that among other things enhance metabolic output. Heterochromatic regions can be under-replicated due to replication fork stalling, a trigger for genome stability in mitotic cycling cells (Nordman et al., 2014).

By comparison, gene amplification is restricted to relatively small segments of the genome that selectively retain the ability to recruit the Origin Recognition Complex (ORC) and MCM2-7 replicative helicase to assemble pre-replicative complexes (pre-RCs) (Austin, Orr-Weaver and Bell, 1999; Claycomb et al. 2002). Whereas DNA replication normally initiates from thousands of sites in chromosomes (origins of replication) during conventional S phases, virtually all replication origins are silenced during locus-specific gene amplification. The few origins that remain active re-initiate DNA replication multiple times to elevate the copy number of neighboring genes. Developmentally programmed gene amplification is a highly regulated process, best illustrated in *Drosophila melanogaster* follicle cells, where several discrete loci are amplified to different levels at different stages of embryonic development (Spradling and Mahowald, 1980; Claycomb et al., 2002). In normal metazoan tissues, endoreplication precedes gene amplification (Calvi et al, 1998 and references therein). This is not the case in cancer cells that amplify protooncogenes or genes that confer drug-resistance (reviewed in Alberton, 2006).

*Tetrahymena thermophila* provides unique opportunities to studying DNA replication and cell cycle checkpoint control. Like all ciliated protozoa, *T. thermophila* harbors two functionally distinct nuclei within its cytoplasm: the transcriptionally silent, diploid ‘germline’ micronucleus and the transcriptionally active, polyploid ‘somatic’ macronucleus (reviewed in Karrer, 2012). ORC-dependent DNA replication is coordinately regulated in vegetative growing Tetrahymena, such that the heterochromatic micronucleus and euchromatic macronucleus replicate their chromosomes at non-overlapping stages of the cell cycle (reviewed in Cole and Sugai, 2012; Zhang et al., 2022). Development is much more complex: multiple rounds of micro- and macronuclear DNA replication, meiosis and mitosis occur in parental and post-zygotic progeny. They all occur within the same cytoplasm in the absence of cell division. Micro- and macronuclear DNA replication are uncoupled. In addition, programmed nuclear death (PND) destroys 3 of the 4 haploid pronuclei and the parental macronucleus (Cole and Soelter, 1997; Yakisich and Kapler, 2004).

The early stages of conjugation (0-8 h) are exclusively devoted to the establishment of germline pronuclei and the transmission of micronuclear chromosomes between mating cells. The genesis of haploid pronuclei for reciprocal exchange between mating partners and production of four genetically identical progeny micronuclei requires five rounds of micronuclear DNA replication. Micronuclei do not resume DNA replication until two post-zygotic nuclei differentiate into macronuclei. Since progeny inhabit the cytoplasm of their parent, the parental macronucleus must be destroyed to confer the phenotype of their progeny.

Macronuclear anlagen formation involves massive genome reorganization. One third of the genome is eliminated by RNA-guided removal of internally eliminated sequences (IESs) (Yao, Fuller and Xi, 2003). Site-specific DNA fragmentation reproducibly converts the 5 micronuclear chromosomes into ~180 discrete macronuclear chromosomes (Eisen et al., 2006). Through endoreplication, macronuclear chromosomes achieve a final copy number of ~45 C (Figure 1A). Furthermore, the single copy 10.3 kb ribosomal DNA locus is excised from its parental chromosome, rearranged into a 21 kb palindromic minichromosome and selectively over-replicated to ~9000 C. Once development is complete, Tetrahymena enters the vegetative phase of its life cycle, where micro- and macronuclear chromosomes replicate once per cell division. Consequently, the integrity of macronuclear chromosomes must be sufficient to minimize genome instability during amitotic vegetative transmission.

**Figure 1.**
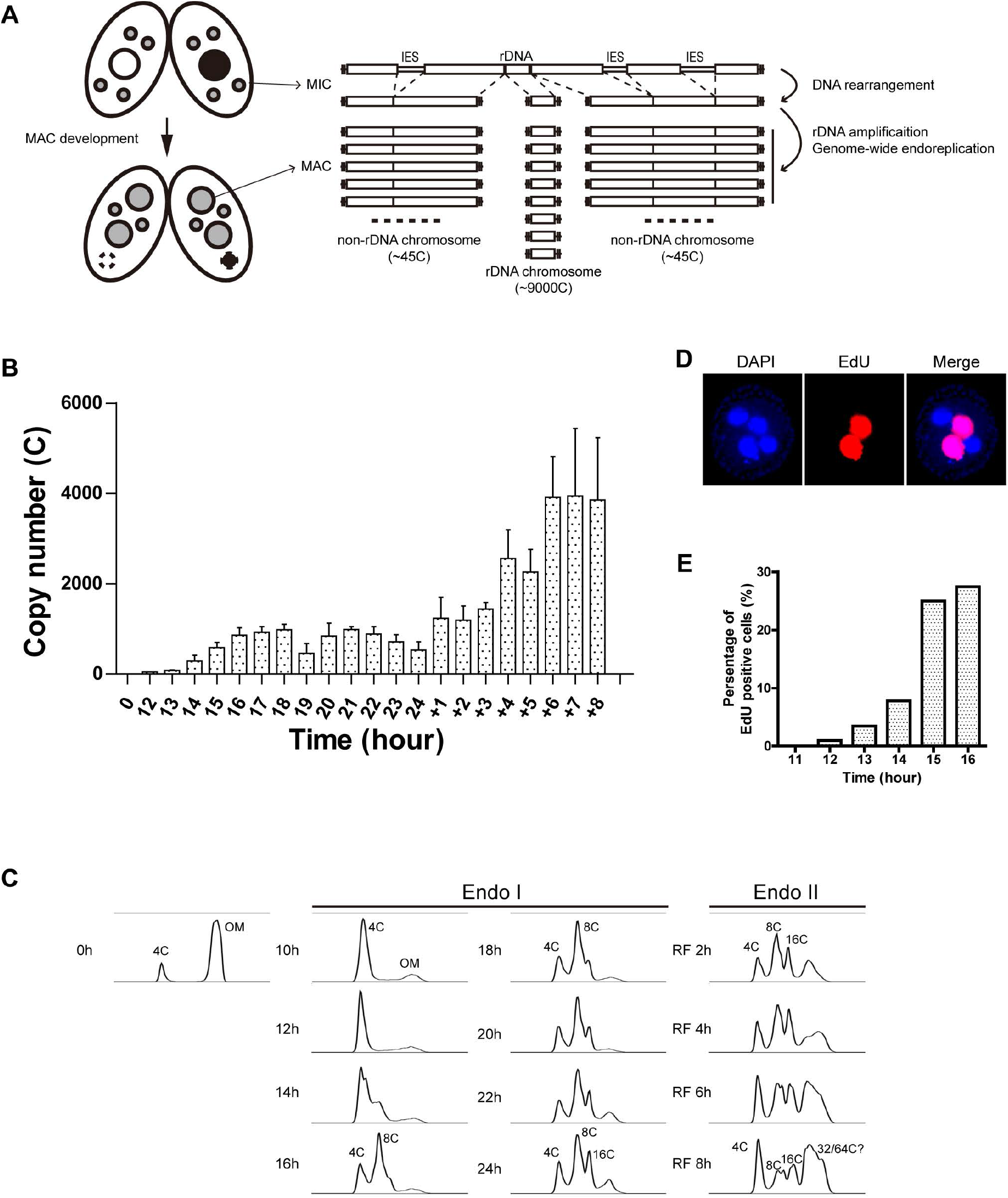
rDNA gene amplification and endoreplication. A. Schematic of mating and subsequent macronuclear development. Upper image: Paired meiotic parental cells of different mating type at an early stage with 4 pronuclei (3 of which will degrade one that will undergo 1 round of DNA replication and nuclear division) and the parental macronucleus. Lower image: exconjugant progeny at a stage with two micronuclei, two developing macronuclei and the degraded parental macronucleus. See text for details on chromosomal changes associated with macronuclear development. B. qPCR analysis on C3 rDNA copy number in the developing macronucleus in a mating between strains SB1934 and SB4202. Samples were collected at 0 h, 12 h-24 h post-mixing. At 24 h, cells were refed, and samples were collected for an additional 8 h. rDNA copy number of each sample was normalized to the sample at 0 h post-mixing (integrated micronuclear rDNA locus). Micronuclear rDNA copy number was used as an internal control for 2-ΔΔCt method. C. Flow cytometry analysis on genomic DNA content during Endoreplication Phase I (mated for 0-24 h) and Endoreplication Phase 2 (mated for 24 h and refed for 1-8 h). OM: old (parental macronucleus). D. A representative image of EdU labeling of anlagen nuclei. Nuclei: DAPI/blue color, active DNA replication: EdU/red. Cells during Endoreplication Phase I have 4 nuclei in each cell. Developing macronuclear anlagen are EdU-positive. E. Quantification of EdU labeled cells as a function of time in mating labeling image showing DNA replication during Endo I.

Recent studies revealed an unexpected relationship between Tetrahymena ORC and the demands for DNA replication. Whereas very modest reductions in Orc1p triggers genome instability in the diploid mitotic micronucleus and polyploid amitotic macronucleus during vegetative growth (Lee et al., 2015), transient reductions of a greater magnitude are readily tolerated in response to DNA replication stress. For example, ORC levels decline 50-fold in hydroxyurea-arrested vegetative cells; however, upon removal of the drug cells immediately enter macronuclear S phase and initiate DNA replication prior to ORC replenishment (Sandoval et al, 2015). ORC-dependent rDNA origins are inactivated in HU-treated cells and replication initiates proximal to the ribosomal RNA promoter during the recovery phase. Furthermore, ORC protein levels naturally fluctuate in mating cells, peaking early in development when four haploid pronuclei are generated and declining during macronuclear anlagen formation, when the demand for DNA replication is the greatest (Lee et al., 2015).

In the work presented here, we determined the relationship between genome-wide endoreplication and locus-specific gene amplification in the developing Tetrahymena macronucleus. We provide evidence that elevated rDNA copy number is generated by two mechanisms rather than one. When ORC levels are high, the rDNA is amplified concurrent with endoreplication of non-rDNA macronuclear chromosomes. When ORC levels decline, the amplification program is shut down, and the rDNA is endoreplicated along with its non-rDNA counterparts. We present quantitative data for the differential regulation of replication initiation and fork elongation in non-rDNA chromosomes as well.

## MATERIALS AND METHODS

### *Tetrahymena* culture and strains

Tetrahymena strains were cultured in 2% PPYS media (2% proteose peptone, 0.2% yeast extract, 0.003% sequestrine) at 30°C. 250 μg/ml penicillin, 100 μg/ml streptomycin and 250 ng/ml amphotericin B (Antibiotic-Antimycotic, Life Technologies). For starvation, Tetrahymena cells were collected from log phase culture, washed twice with 10 mM Tris-HCl (pH 7.4), and starved in the same buffer supplemented with 250 μg/ml penicillin, 100 μg/ml streptomycin and 250 ng/ml amphotericin B for 18 h. For mating, Tetrahymena cells were starved 18 as described, mixed with equal number at the final cell density of 2.5×10^5^cell/ml and then incubated under stationary conditions at 30° C. All Tetrahymena strains used in this study are listed in Table 1.

**Table 1.** *T. thermophila* strains used in this study.

| **strain** | **Micronuclear genotype** | **Macronuclear phenotype** |
| --- | --- | --- |
| CU427 | *chx1-1/chx1-1* | paromomycin-sensitive<br>cycloheximide-sensitive |
| CU428 | *mpr1-1/mpr1-1* | paromomycin-sensitive<br>6-methylpurine-sensitive |
| SB1934 | rDNA [C3-1]/rDNA [C3-1];<br>*mpr1-1/mpr1-1* | paromomycin-sensitive<br>6-methylpurine-sensitive<br>rDNA [B] |
| SB4202 | *pmr1*[C3-1]/*pmr1*[C3-1];<br>*CHX1*[C3]/*CHX1*[C3] | paromomycin-sensitive<br>cycloheximide-resistant<br>rDNA [B] |
| g-H3.2+GFP-3 | *hht2*[3'neo2, GFPc]<br>*hht2*[3'neo2, GFPc] | paromomycin-sensitive |

### 5-ethynyl-2’-deoxyuridine (EdU) labeling

Click-iT EdU Alexa Fluor 594 Imaging Kit (Life Technologies) was used to identify cells undergoing DNA replication using the EdU labeling assay. Basically, EdU was added to the mating culture at the final concentration of 100 μM and mating cells were labeled at 30°C for 20 min. The cells were then washed, fixed, and reacted with Click-It reagent according to manufacturer’s instructions (Life Technologies). DAPI was used to stain the nuclei. Both DAPI staining and EdU labeling were detected by fluorescence microscopy.

### Quantitative PCR (qPCR)

Genomic DNA was isolated from Tetrahymena cells according to a previously described protocol (Zhang et al., 1997). qPCR was performed in 96 well plates on the StepOnePlus Real-Time PCR Systems (Applied Biosystems). qPCR was done in a 10 μl reaction including 5 μl Power SYBR^®^ Green PCR Master Mix (Applied Biosystems), with 1-5 ng genomic DNA as a template, and 200 nM forward and reverse primers. Each DNA sample was assayed in triplicate. Water was used to replace genomic DNA for the negative control.

### Flow cytometry

DNA content of micronuclei and anlagen macronuclei of mating cells was quantified by flow cytometry (Sandoval et al. 2015). 5×10^5^ cells from mating cultures were collected by centrifugation at 3,000 rpm for 3 min. Cells were washed once using 10 mM Tris-HCl (pH7.4) and re-pelleted by centrifugation. 0.5 ml of TMS buffer (10mM Tris-HCl [pH 7.4], 10 mM MgCl_2_, 3 mM CaCl_2_, 0.25 M sucrose, 0.2% NP-40) was added to lyse cells under condition that keep nuclei intact. Propidium iodide (PI) and RNase A were then added to the samples at a final concentration of 20 μg/ml and 200 μg/ml, and samples were incubated in the dark at RT for 30 min. Samples were analyzed on Becton Dickinson FACSAria II flow cytometer (BD Biosciences). A total of twenty thousand events were collected for each sample for data analysis. Data analyses were performed using Flowjo version 10 software. The number of events versus the relative DNA content was plotted in histograms.

### DNA fiber analysis

DNA fiber analysis was performed as previously described (Lee et al, 2015). Tetrahymena cells were pulse-labeled with 400 μM IdU (Sigma) at 30°C for 10 min. Then cells were washed once with 1×PBS. Cells were resuspended in pre-warmed fresh media (for vegetative cells) or 10 mM Tris-HCl (pH7.4) (for mating cells) with 100 μM CldU (MP Biomedicals) and labeled for 10 min. After two washes with PBS, the cell density was adjusted to 1×10^6^cell/ml. Preparation and immunostaining of DNA fibers were performed as previously described. (Chastain et al, 2006; Schwab & Niedzwiedz, 2011; Stewart et al, 2012) with the following modifications. Briefly, after fixation and HCl treatment, slides were washed three times with 1×PBS, and 5% BSA in PBS was used to block slides for 30 min. Mouse α-BrdU (1:50, Becton Dickson) which recognizes IdU and rat α-BrdU (1:100, Accurate Chemical) which recognizes CldU in 5% BSA were then added onto slides. After 1 h incubation, the slides wash washed three times with 1×PBS and incubated for 30 min with secondary antibodies: Alexa Fluor 568 goat anti-mouse IgG (1: 100, Invitrogen/Molecular probes) and Alexa Fluor 488 goat anti-rat IgG (1: 100, Invitrogen/Molecular probes). Next, slides were washed three times with 1×PBS, dehydrated with an ethanol series and mounted with SlowFade Gold antifade (Invitrogen). During immunostaining, all antibodies were diluted in 5% BSA in 1x PBS, all incubations were performed at 37°C, and all wash steps were done at RT.

DNA fiber images were taken by a Nikon A1R+ confocal microscope with 600x magnification. Measurements of track length were performed with Nikon NIS-Elements software. Inter-origin distance was defined as the distance between the centers of two red segments in either green-red-green-red-green or green-red-gap-red-green tracks. Fork velocity was determined by measuring the length of the green segment in a red-green track or red segments in a green-red-gap-red green track. GraphPad Prism software was used to analyze the statistical significance, and p-values were determined by Mann-Whitney test for single comparisons or Kruskal-Wallis test for multiple comparisons.

### Two-dimensional (2D) gel electrophoresis of DNA replication intermediates (RIs)

Total genomic DNA was prepared as previously described (Zhang et al., 1997). 200 μg of genomic DNA were digested with restriction enzymes at 37° C for 4 h, precipitated by ethanol, and resuspended in 400 μl of TNE buffer (100 mM NaCl, 10 mM Tris, 1 mM EDTA, pH 8.0). The digested DNA was then applied to 200 μl of benzoylated naphthoylated DEAE (BND)- cellulose (Sigma-Aldrich). RIs was bound to BND-cellulose, and non-specific binding DNA fragments were washed out with 400 μl of TNE buffer for five times. 200 μl of 1.8% caffeine in TNE was used to elute RIs from BND-cellulose. DNA was then precipitated with isopropanol, washed with 70% ethanol and resuspended in TE buffer. For B rDNA strains, the enriched RIs were directly applied onto gel electrophoresis. For strains containing C3 rDNA in the developing MAC and B rDNA in the parental MAC, RIs were further digested by SphI to separate parental B rDNA from progeny C3 rDNA 5’ NTS fragments.

Neutral-neutral 2D gel electrophoresis was performed according to previous description (Zhang et al. 1997). Typically, 5-10 μg of BND cellulose-enriched RIs were loaded on a 0.4% agarose gel, and the first dimensional gel was run in 1xTAE buffer at 1.5 V/cm for 18 h at RT. The gel was visualized by ethidium bromide staining, and gel slice from each lane was cut with correct size range of RIs and rotated 90 degree for second dimension gel electrophoresis. Gel slices were inserted into 1% agarose gel, and the second dimensional gel electrophoresis was performed in 1xTBE buffer (89 mM Tris, 89 mM boric acid, 2 mM EDTA, pH 8.0) containing 0.5 μg/ml ethidium bromide at 3 V/cm for 18 h at 4°C. Southern blot analysis was carried out to detect the patterns of RIs as previously described (Zhang et al. 1997).

## RESULTS

### Quantitative analysis of rDNA gene amplification in the developing macronucleus

All eukaryotes contain multiple copies of ribosomal RNA genes to meet the high demands for protein synthesis. Virtually all species encode large tandem rDNA repeat arrays in one or more chromosome (reviewed in Iida and Kobayashi, 2019). In Tetrahymena this is achieved by the developmentally programmed excision of the single copy rRNA gene, formation of a palindromic 21 kb minichromosome, and re-replication to a final copy number ~9000 C. Meanwhile, the remainder of the genome is endoreplicated to a final copy number of 45 C. To obtain new insights into the production of rDNA minichromosomes, we used quantitative PCR (q-PCR) to examine the temporal increase in rDNA and non-rDNA copy number during development. Persistence of parental macronucleus (PM) in progeny until late stages of macronuclear development obscured our ability to assess de novo synthesis of rDNA minichromosomes. To circumvent this problem for the rDNA, we mated heterokaryon strains SB1934 and SB2402, both of which are homozygous for the C3 rDNA allele in the diploid germline micronucleus, but contain B rDNA in the polyploid somatic macronucleus (Table 1). C3-specific primers were used to quantify the increase in abundance of rDNA minichromosomes in developing macronuclei, while a second primer set amplified the single integrated rDNA gene in the micronucleus. The 2^-ΔΔCt^ method was used for quantification of macronuclear rDNA copy number (Livak & Schmittgen, 2001). Genome-wide endoreplication was assessed by flow cytometry.

A wave of rDNA synthesis was detected in the developing macronucleus between 13-16 h (from 4 C to ~800 C), followed by a plateau in starved mating cells (Figure 1B, linear graph plot). One round of endoreplication was achieved at this time (Figure 1C, 8 C PI peak). Whereas the rDNA copy number did not increase further in starved mating cells (18 h to 24 h), a significant subpopulation completed another round of endoreplication by 24 h (Figure 1C). rDNA replication resumed upon re-feeding of mated cells at 24 h, achieving a copy number of >4000 C by 8 h after re-feeding (RF) (Figure 1B). The majority of re-fed mating cells achieved a ploidy of 32-64 C (Figure 1C, RF 1-8 h). The periodicity of PI flow cytometry peaks is consistent with previous Feulgen staining analyses of individual cells (Li et al., 2006). These oscillations are expected for genome-wide endoreplication cycles (Gap-S-Gap-S). They are not consistent with continuous re-replication in a single, extended macronuclear S phase.

To assess genome-wide endoreplication at the cellular level, the nucleoside analog of thymidine, 5-ethynyl-2’-deoxyuridine (EdU) was used to pulse-label cells to measure active DNA synthesis. Cells were collected at hourly intervals, fixed and subjected to click chemistry for microscopic examination or flow cytometry (to measure bulk DNA content in the population). EdU labeling was detectable in macronuclear anlagen at 14 h post-mixing (Figure 1D), near the onset of rDNA gene amplification (Figure 1C). EdU incorporation increased dramatically during the 15 h and 16 h pulses (Figure 1E). Flow cytometry analysis data showed a similar pattern, the change in anlagen DNA content was evident at 14 h and increased significantly at 15 h to 16 h (Supplemental Figure 1). ~30% of mating cells were EdU positive following a 20 min pulse label during the 4 C to 8 C endoreplication window, consistent with endocycling within with an S phase of ~40 min (Figure 1E, 18 h). The 2 h Gap-S-Gap-S endocycle in starved mating populations was asynchronous, as anticipated based on differences in the timing for mating pair formation. By comparison, macronuclear S phase constitutes about 1/3 of the ~3 h vegetative cell cycle (Zhang et al., 2022).

### Kinetics of rDNA and non-rDNA copy number increases during development

A limitation of our initial approach is that rDNA and non-rDNA abundance were assessed by different methods. To overcome this we initiated a cross to assess rDNA and non-rDNA copy number changes by the same method. A C3 rDNA heterokaryon strain (SB1934: C3 rDNA micronucleus, B rDNA macronucleus) was mated to a histone H3 heterokaryon strain (g-H3.2+GFP-3: gfp-histone H3 micronucleus; wildtype histone H3 macronucleus), in which the coding sequence of the micronuclear HHT2 gene was fused to green fluorescent protein (GFP) (Table 1; Supplemental Figure 2). Allele-specific q-PCR was used to exclusively monitor C3 rDNA and HHT2-GFP copy number in the developing macronucleus of progeny. Four important pieces of information were obtained. First, rDNA amplification initiated prior to endoreplication of the non-rDNA HHT2 locus (Figure 2A, T=12 h). Second, in contrast to all other developmental systems, gene amplification and endoreplication occur concurrently. Unexpectedly, rDNA amplification was restricted to a relatively short developmental window in starved mating cells (Figure 2A; T=10 h-16 h). Third, within the limits of resolution, there appears to be a limiting factor or environmental cue that concurrently down regulates both DNA replication programs (Figure 2A; transition point: T=16 h). Finally, upon re-feeding, mated cells resume replication of rDNA and non-rDNA at chromosomes at comparable rates. We conclude that rDNA replication initiation is subjected to multiple levels of regulation: cell cycle control during vegetative S phase, gene amplification during early macronuclear development, and endoreplication during the later stages of macronuclear development. As previously reported, the increased demand for DNA replication during macronuclear development in inversely correlated to the abundance of ORC (Lee et al., 2015). Gene amplification accounts for ~20% of the final rDNA copy number and endoreplication generates the rest.

**Figure 2.**
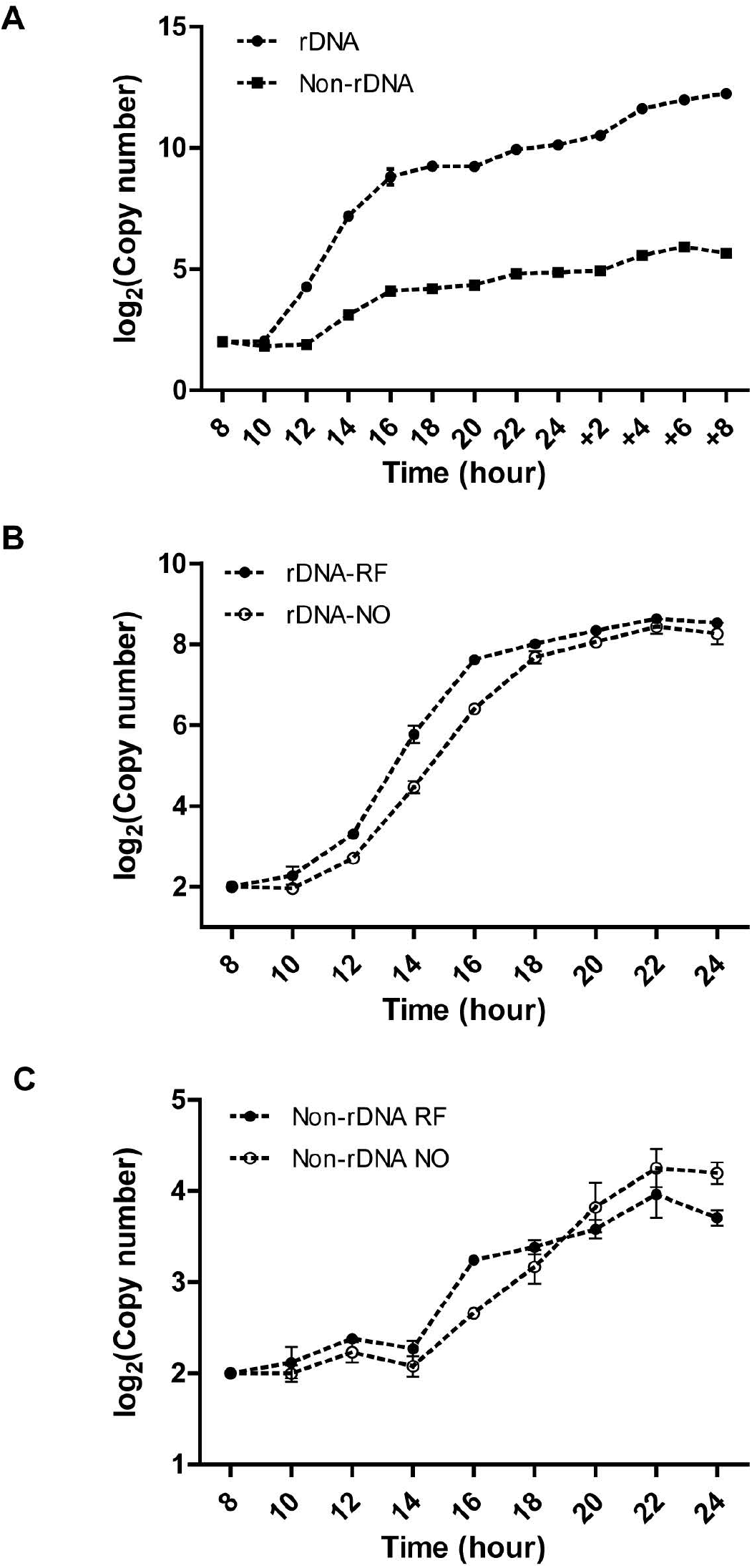
Q-PCR analysis of rDNA and non-rDNA (H3-GFP) chromosome copy number changes during *macronuclear* development. Heterokaryon strains SB1934 and g-H3.2+GFP-3 were mated. Q-PCR analysis was performed with C3 rDNA and H3-gfp locus-specific primer sets to quantify DNA copy number in the newly developing macronucleus. The copy number of both rDNA and non-rDNA at 0 h was set to 2 C. (A) Log_2_ scale of copy number changes of rDNA and non-rDNA chromosomes in mated Tetrahymena that were refed at 24 h post mixing. (B) Log_2_ scale of copy number change of rDNA copy number in mated Tetrahymena that were refed at 10 h versus 24 h post-mixing. RF: early refed at 10 h; NO: normal refed at 24 h. (C) Log_2_ scale of copy number change of non-DNA (H3-gfp locus) copy number in mated Tetrahymena that were refed at 10 h or 24 h post-mixing. RF: early refed at 10 h; NO: normal refed at 24 h.

### Effect of re-feeding on endoreplication phases 1 and 2

One possible explanation for the plateau in rDNA and non-rDNA replication in starved mating cells (Figure 1B and 1C, Figure 2A) is that re-feeding is required to generate new proteins and/or DNA precursors to sustain further rounds of DNA replication. One prediction is that early re-feeding might eliminate the starvation plateau and possibly support more extensive rDNA gene amplification as well. To explore these possibilities, we compared mating cells that were re-fed early (10 h post-mixing) to our normal regiment (refeeding at 24 h). Subtle differences were detected. For example, rDNA amplification was modestly accelerated in early re-fed mating cells, but there was no net increase in copy number (Figure 2B). The amplification ‘window’ was neither shortened nor lengthened: rDNA copy number plateaued in the 18-24 h interval, regardless of whether mating cells were re-fed. Early re-feeding had no obvious impact on the endoreplication program as well, as assessed through copy number analysis of the HHT2-gfp allele (Figure 2C). The collective data indicate that there are other underlying factors that control the temporal/developmentally program for overreplication of rDNA and non-rDNA chromosomes.

### Replication initiation and elongation in endoreplicating non-rDNA chromosomes

In previously published work we discovered that ORC protein levels are dynamically regulated during development (Lee et al., 2015). The observed changes are counterintuitive for ORC-driven replication initiation. During the period for meiotic and post-zygotic micronuclear DNA replication (0-9 h), Orc1p level are elevated 3-fold relative to vegetative G1/S cultures, despite an ~25-fold reduction in the amount of DNA synthesis. Orc1p levels subsequently decline ~30-fold (0.1X vegetative level), reaching a minimum during endocycle phase II, when the developmental DNA replication load is the greatest (Lee et al., 2015). Mcm6 protein levels correlate well with Orc1p during development and are similarly diminished during vegetative S phase in ORC1 knockdown mutants, suggesting that the abundance of these pre-RC components is coordinately regulated. Two studies, one in Drosophila and another in

To examine DNA replication in the developing macronucleus, we employed DNA fiber imaging to measure origin activity and fork elongation rates in endoreplicating non-rDNA chromosomes relative to vegetative DNA synthesis. Our data revealed a 12% reduction in the average inter-origin distance in endocycling cells (vegetative S phase: 26.9 kb; endoreplication phase I: 24.2 kb; endoreplication phase 2: (early) 23.8 kb and (late) 24.5 kb (Figure 3A). Hence, more initiation events occur in endoreplicating macronuclei, despite the decreased abundance of ORC and MCM proteins. Consistent with reduction in the replicative helicase, we observed a >25% decrease in the rate of replication fork elongation (Figure 3B; vegetative S phase, 0.87 kb/min; endoreplication phase I, 0.77 kb/min; endocycle phase 2 (early and late) 0.65 kb/min. These data are indicative of compensatory changes in replication initiation and elongation in endocycling cell populations.

**Figure 3.**
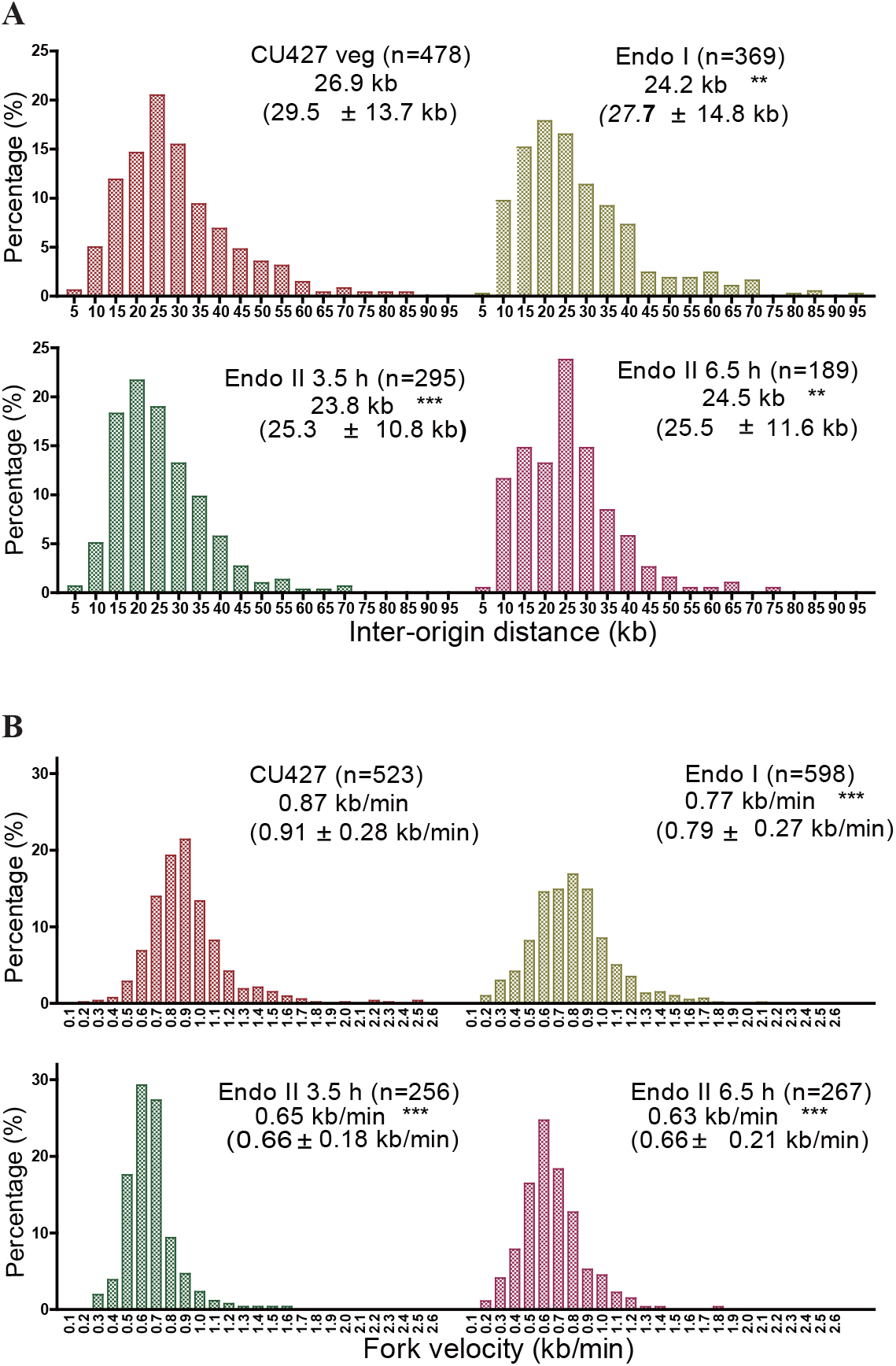
DNA fiber analysis on endoreplicating macronuclear anlagen. Mated cell progeny from a CU428 x CU427 cross were collected at different times and process for DNA fiber analysis. Endo I: mate cell progeny were pulse with CldU and chased with IdU at 15.5 h post-mixing. Endo II: mated cell progeny were refed at 24 h and sequentially labeled with CldU and IdU at 3.5 h or 6.5 h after refeeding. A log phase CU427 vegetative culture was used as a control. A. Inter-origin distance (IOD) comparison between log phase and endoreplicating cells. B. Fork velocity comparison between log phase and endoreplicating cells. The frequency distribution of DNA fibers with different IODs or fork velocities were graphed. For each group, the median IOD or fork velocity was calculated. The mean with S.D. is indicated in parenthesis. N: the total number of DNA fibers scored. Statistical significance was determined using the Kruskal-Wallis test followed by Dunn’s test. Each sample from endoreplicating cells was compared to log phase CU427. **, p < 0.01; ***, p < 0.001.

### Localization of aberrant replication intermediates in endoreplicating rDNA minichromosomes

Two-dimensional (2D) gel electrophoresis of rDNA replication intermediates (RIs) previously demonstrated that the origins that amplify the rDNA in the developing macronucleus also control DNA replicate during vegetative cell cycles (Figure 4A schematic) (Zhi et al., 1998; MacAlpine et al., 1998). In vegetative S phase replication initiates exclusively from origins in the 5’ non-transcribed spacer (NTS), as evidenced by the presence of bubble-to-Y arc RIs, and absence of complete simple Y arcs in HindIII-digested DNA (Figure 4A, restriction map; Figure 4B, upper left panel). Mutations in 26T RNA, a unique integral ORC subunit, inhibit ORC binding to origin-proximal type I elements and block initiation from these 5’ NTS origins (Mohammad et al. 2007; Donti et al., 2009). We more recently identified unusual replication intermediates (RIs) that form in wild type Tetrahymena during endoreplication phase 2 in re-fed mating cells (Figure 4B and 4C (HindIII), middle panel) (Lee et al., 2015). Aberrantly migrating simple Y arcs suggest that the known 5’ NTS origins are passively replicated in a subpopulation of molecules. The aberrant Y arcs pattern is consistent with extensive DNA unwinding and/or accumulation of persistent RNA:DNA hybrids, similar to what has been reported in *S. cerevisiae* senetaxin mutants (Alzu et al, 2012). The aberrant RIs (Figure 4C, (HindIII, right panel) and their sensitivity to mung-bean nuclease were reported in the vegetative *Tetrahymena* TXR1 knockout strain, defective in histone H3K27 monomethylation (Gao et al, 2013; Lee et al, 2015).

**Figure 4.**
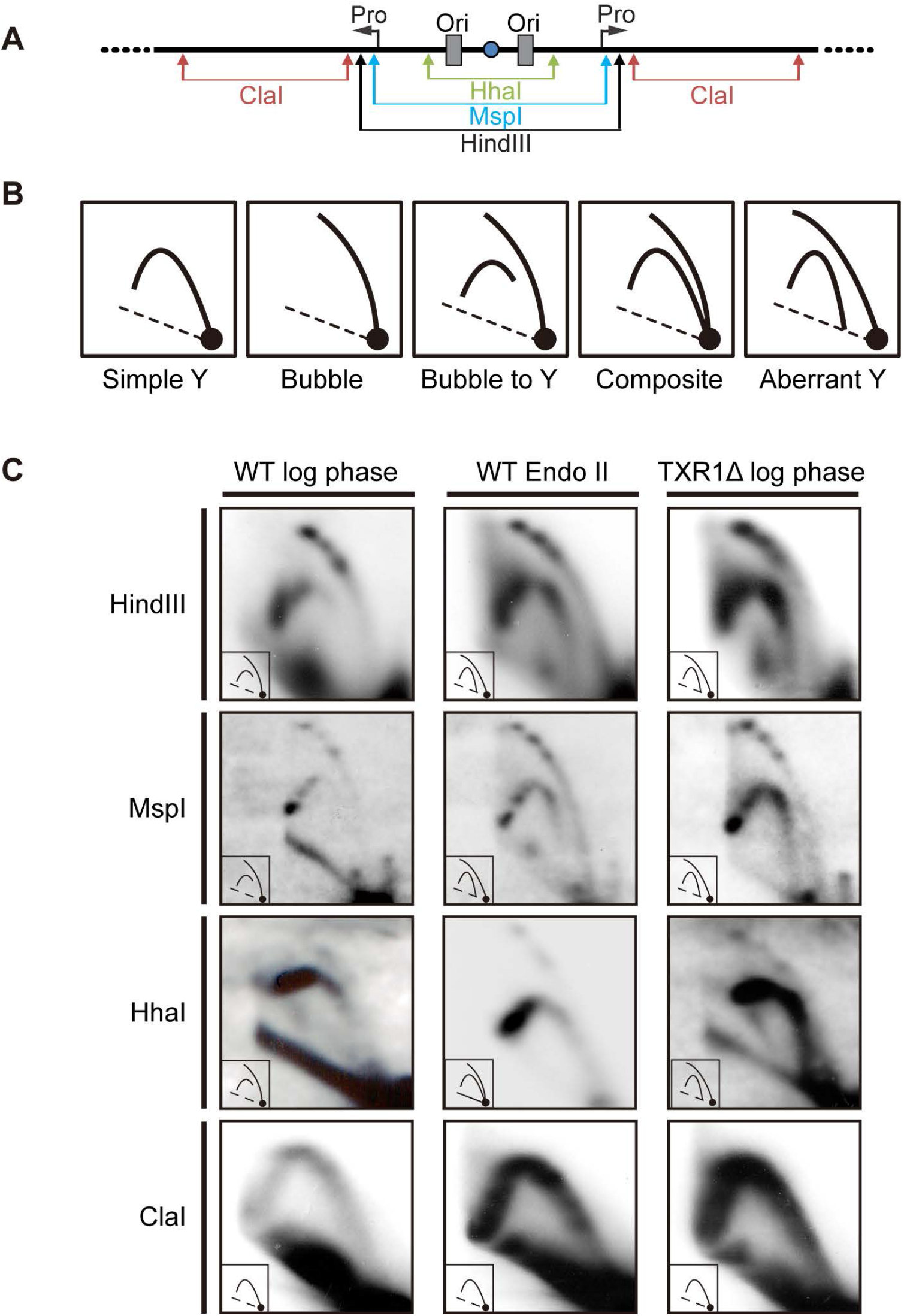
Two-dimensional gel analysis of rDNA replication intermediates (RIs) in endoreplicating macronuclei. A. Schematic of the rDNA 5’NTS fragments generated by restriction digestion for 2D gel analysis with rDNA 5’ NTS or coding region probes. H3, Hind *III* sites is at nucleotide (nt) position 2132 in the palindromic 21 kb rDNA minichromosome; MspI: nt 1906; HhaI: nt 1347; ClaI: nt 2169-7621. The two origins (ORI) resides within the tandem 430 bp duplication designated as Domain I and Domain II (ORC binding sites: type IA and type IB elements (nt 762-794 and nt 1193-1224), Pro: 35S rRNA precursor start site (nt 1887) of the rDNA minichromosome. B. Schematic of normal and aberrant DNA replication intermediate patterns. Different types of RIs may be detected by 2D gel analysis. Simple Y arc: replication from outside the examined DNA interval (passive replication). Bubble arc or composite bubble-to-Y arc pattern: replication initiation within the examined region. Composite with complete bubble and Y arcs: both active and passive replication within the examined region. Aberrant Y: replication intermediate that fails to intersect the 1N spot corresponding to unreplicated duplex DNA. Diagonal dashed line, migration of bulk linear duplex DNA fragments. C. 2D gel analysis of wild type log phase vegetative cells (CU428), endoreplicating mated cells (SB1934 x SB4204 cross, mated 24 h and refed for 6 h) and a vegetative log phase TXR1 knockout mutant (generated by macronuclear gene disruption and phenotypic assortment) (Gao et al., 2013).

In an effort to localize the region responsible for the aberrant RIs, we performed 2D gel analysis on DNA digested with different restriction enzymes (Figure 4A). The vegetative TXR1 mutant was used as a reference point for comparative analysis to wild type vegetative and endocycling RIs. We specifically set out to determine whether the rDNA promoter region (responsible for rRNA biogenesis) or upstream non-coding sequences were involved. MspI cleaves 18 bp downstream of the rRNA start site, and HhaI digests just downstream of the tandem ORC binding sites (ORI). Aberrant simple Y arcs were detected in the MspI 5’ NTS fragment, but not in the HhaI-digested sample (Figure 4B, simple Y vs. aberrant Y; Figure 4C). Furthermore, normal migrating simple Y arcs were detected in the ClaI coding region fragment. The close proximity of the MspI site to the rRNA start site (18 bp) in conjunction with normal migrating ClaI coding region RIs, suggest that stalled 35S rRNA transcripts are not responsible aberrant RI production. The collective data suggest that rDNA replication initiates downstream of the nucleosome-free 5’ NTS origins that are bound by Watson:Crick RNA-DNA base pairing to ORC (Mohammad et al. 2007), and upstream of the rRNA promoter (Pan and Blackburn, 1995) in a large fraction of replicating molecules. This interval is comprised of repetitive DNA sequences, the type II a-m elements. The implications are discussed below.

## Discussion

In conventional mitotic cell cycles (G1-S-G2-M) DNA content oscillates by a factor of two. Chromosomes duplicate once in S phase and are transmitted equally to daughter cells. Developmentally programmed endoreplication (Gap-S-Gap-S) increases DNA content on a genome-wide scale, enhancing a cell’s capacity to increase in size and/or perform specialized functions. For example, polyploid Drosophila salivary glands secrete copious amounts of proteins required for pupation (Berendes, 1965). Polyploid mammalian placental trophoblast giant cells provide a barrier between maternal and fetal blood supplies, and secrete hormones that regulate maternal blood flow (reviewed in Cross, 2005). Although locus-specific gene amplification is restricted to small segments of chromosomes, it achieves a similar objective, illustrated by the amplified Drosophila chorion genes which direct the production of essential eggshell proteins that encase the developing embryo (Spradling and Mahowald, 1980).

Ciliated protozoa provide unique opportunities to study these and other unconventional DNA replication programs. This ancient lineage contains large protozoan species, some of which are visible to the naked eye. In the case of *Stentor coeruleus*, single cell sequencing demonstrated that chromosome copy number scales with the size of individual organisms (Slabodnick et al., 2017). On average, the rDNA contig is present in over 1M copies per organism/cell and non-rDNA macronuclear chromosomes achieve a copy number of 20,000-60,000.

The partitioning of germline and somatic functions into diploid, heterochromatic micronuclei and polyploid, euchromatic macronuclei, respectfully, is a defining feature of the early branching Ciliophora phylum. Synchronized mass mating has been exploited to discover and study molecular events and biochemical pathways for genome reorganization in the developing macronucleus. In the work presented here, we mated *Tetrahymena thermophila* heterokaryons to study endoreplication and gene amplification. The copy number of marked rDNA and non-rDNA chromosomes was determined in progeny macronuclei, including early timepoints in development, when the parental macronucleus was still present. These studies revealed a more complex DNA replication program than previously realized.

Endoreplication in Tetrahymyena occurs in two distinct phases, separated by a prolonged gap phase that is experimentally terminated by re-feeding (Figures 1 and 2). However, early refeeding is not sufficient to override cell cycle arrest of endoreplication phase I at the 8 C stage (Figure 2). We proposed that nutrients and the activation of nutrient signaling pathways are required for endoreplication phase 2, as reported for TOR and SNAIL signaling in Drosophila (Zeng et al., 2020), but that they only function when intrinsic cellular programs are prepared to respond. Some possible internal drivers for the transition to endoreplication phase 2 include programmed DNA rearrangement (reviewed in Noto and Mochizuki, 2017), genome-wide conversion of heterochromatin into euchromatin (reviewed in Chalker 2008; Gao et al., 2013), new zygotic gene expression, modulation of the ASI2 signal transduction pathway, which is required for initiation of the endoreplication program (Li et al., 2006; Yin, Gater and Karrer 2010), and degradation of the parental macronucleus (Akematsu, Pearlman and Endoh, 2010). Extrapolating to mating in the wild, which occurs in freshwater ponds, nutrient deprivation serves as the environmental signal for seeking out a mating partner and initiating the conjugation program (reviewed in Cole and Sugai, 2012). According to our data, an influx in nutrients would be required to complete macronuclear development.

A major distinction of ciliates is that endoreplication and gene amplification are not associated with terminal differentiation. The products of these alternative DNA replication programs are propagated during ‘vegetative’ cell cycles, when macronuclear chromosomes replicate once and only once during S phase (Figure 5). With the exception of Drosophila adult rectal papillae cells that have a limited, error-prone proliferative lifespan (Fox, Gall and Spradling, 2010), developmentally programmed endoreplication and gene amplification in metazoa occur in terminal differentiated cells. Chromosome integrity can be relaxed as chromosome do not need to be partitioned to daughter cells (Hassel et al., 2013; Mehrotra et al 2008). This situation is illustrated in the Drosophila salivary gland endocycle where cytologically visible segments of the genome are under-replicated (Nordman et al., 2014; Yorch and Spradling, 2014). Furthermore, ‘onionskin’ DNA structures/stalled forks generated during amplification of Drosophila chorion gene loci are tolerated since the affected chromosomes are not transmitted to daughter cells (Osheim and Miller, 1988; Claycomb et al., 2002). When both occur, they are sequentially ordered-endoreplication precedes locus-specific gene amplification (Figure 5).

**Figure 5.**
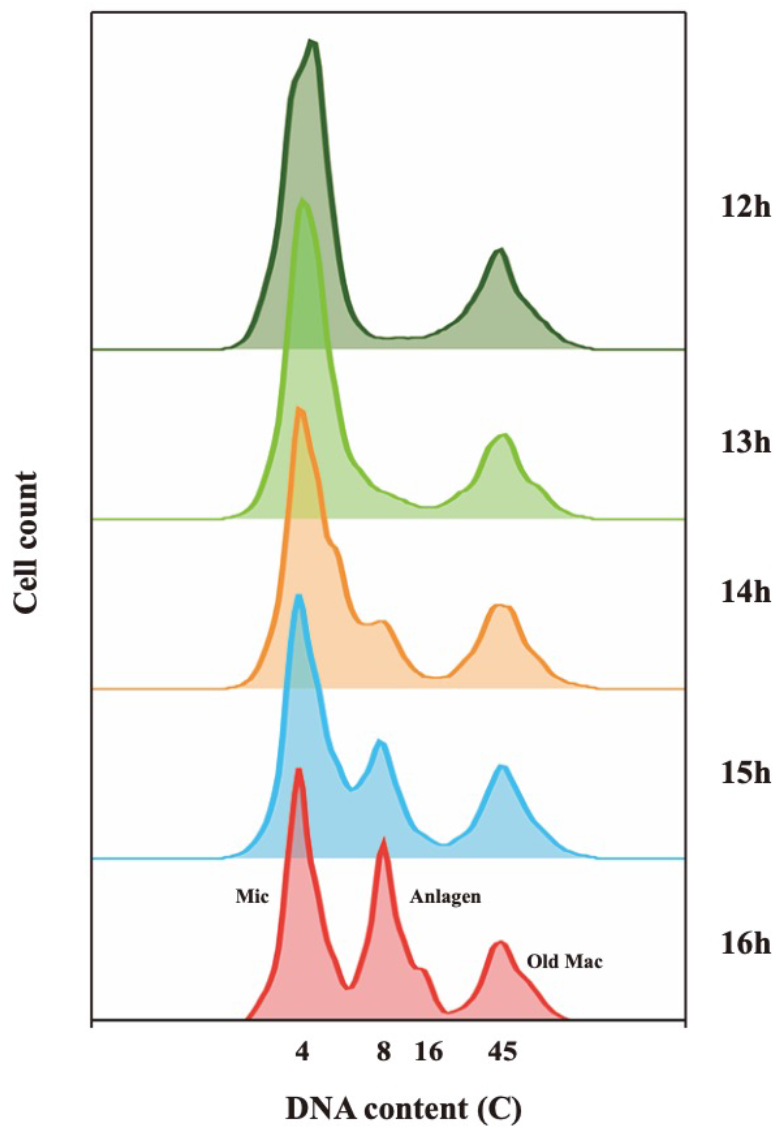
Comparative profiling of endoreplication and gene amplification developmental programs in metazoa and Tetrahymena. Upper panel-sequential DNA replication programs in terminally differentiated cells in flies and humans. Lower panel-sequential DNA replication programs in the developing macronucleus and subsequent entry into the amitotic vegetative cell cycle.

The only known Tetrahymena gene devoted to endoreplication is ASI2, a putative transmembrane signal transduction protein that is required to sustain early and late rounds of endoreplication in the developing macronucleus (Yin, Gater and Karrer, 2010). For comparison, Notch signaling plays a critical role in the mitosis-to-endocycle transition in Drosophila in follicle cells (Sun et al., 2008). Perturbations of intrinsic signaling by 14-3-3 gamma, an inhibitor of the G2-to-M transition, or the local tumor environment promote endoreplication in human cancer cells (Gomes et al., 2017; Tan et al, 2018). DNA double strand breaks (DSBs) trigger endoreplication in normal *Arabidopsis thaliana* endosperm and are thought to inhibit the G2-to-M transition (Adachi et al., 2011). Like metazoan, DNA replication in macronuclear anlagen involves a switch from a mitotic cell cycle, albeit an altered one, in which micronuclear DNA replication and mitosis (nuclear division, without cytokinesis) precedes endoreplication phase 1. Remarkably, positional information dictates whether a nucleus differentiate or exits the cell cycle as a micronuclei (Figure 1D).

Early in macronuclear development, sequence-specific double strand breakage occurs to fragment the 5 germline chromosomes into 180 macronuclear chromosomes. An RNA-guide mechanism eliminates over 5000 internal DNA segments by breakage and joining reactions. It is plausible that DSB formation events trigger the mitosis-to-endocycle transition in Tetrahymena, similar to Arabidopsis endosperm. Maternal expression of PDD1, the nucleating factor for programmed DNA elimination, is required for endoreplication (Coyne et al, 1999) and PDD1-associated proteins are required for ligation of DSBs that among other things removes repetitive DNA sequences, including retrotransposons (Xu et al., 2015). The precision of these processes appears to be adequate to create intact chromosomes for subsequent propagation during the vegetative stage of the life cycle.

Existing data suggest that rDNA amplification and genome-wide endoreplication do not induce genome instability in Tetrahymena. Consequently, aberrant DNA structures or collapsed replication forks are either not generated, are subsequently resolved, or are simply tolerated. Our 2D gel data clearly demonstrate the production of aberrant replication intermediates (Figure 4). Activation of the ATR intra-S phase checkpoint is consistent with DNA damage/fork arrest resolution. The third possibility-toleration-is plausible based on prior studies on vegetative knockdown mutations of ORC1 and TIF1, both of which activate the ATR intra-S phase checkpoint response, but do not arrest cell cycle progression (Yakisich et al. 2006; Lee et al., 2015). Lagging macronuclear chromosomes are observed in these mutants, slowing vegetative cell cycle progression. These strains can be propagated indefinitely in the vegetative phase of the life cycle. However, their diploid micronucleus is genetically unstable, producing DNA deletions and rearrangements, and even entire the loss of entire chromosome that result in sterility.

Four features of macronuclear chromosomes may factor into Tetrahymena’s ability to continuously propagate endoreplicated and amplified chromosomes. First, macronuclear chromosomes are pure euchromatin and there is no evidence for early and late macronuclear DNA replication programs (L Zhang and GM Kapler, unpublished results). With the exception of a few rare minichromosomes that are only transiently propagated during early vegetative growth, there are no known regions that block fork elongation and render macronuclear chromosome sensitive to chromosome loss (Feng et al., 2017). Progeny can propagate indefinitely in the vegetative state. By analogy, endoreplicating Drosophila salivary gland chromosomes undergo fork collapse at many sites (Yarosh and Spradling 2014, Nordman et al, 2014). Genome instability severely limits proliferative lifespan of the rare cell type that can propagate after endoreplication-Drosophila rectal papillae (Fox et al, 2010). Second, in contrast to amplified regions in Drosophila and Sciara chromosomes (Liang et al, 1993; Spradling and Mahowald, 1980), the Tetrahymena rDNA locus resides on a small autonomous minichromosome. The demands for bidirectional fork progression are minimal, since diverging forks only need to traverse only 10-12 kb to propagate the entire rDNA minichromosome (Zhang et al. 1997). To the best of our knowledge, incomplete DNA replication and fork collapse are not developmentally programmed in Tetrahymena. In contrast, stalled/collapsed forks generate during multiple rounds of chorion gene amplification produces onionskin chromosomes that cannot be partitioned to daughter cells (Osheim, Miller and Beyer, 1988). Third, all macronuclear chromosomes lack centromeres and segregate by amitosis. Consequently, anaphase bridges do not form and breakage-bridge-fusion (BFB) cycles are averted. Whereas lagging macronuclear chromosomes are generated at a low frequency in wild type cell, they are inconsequential. In contrast, BFB cycles in mitotic germline micronuclei render pronuclei inviable (Yakisich et al., 2006; Lee et al. 2015). Finally, a ‘self-correcting’ DNA copy number control mechanism somehow maintains genic balance of macronuclear chromosomes (Doerder et al, 1981; Eisen et al., 2006). Aberrant chromosomes, if generated may be eliminated through a surveillance mechanism, analogous to programmed DNA elimination in the early developing macronucleus (reviewed in Noto and Mochizuki, 2017). Despite the existence of a robust intra-S phase DNA damage response, checkpoint activation does not result in terminal cell cycle arrest and activation of the apoptotic program (Sandoval et al., 2015).

As anticipated from previous work, our experiments showed that endoreplication does not precede gene amplification. Both DNA replication programs occur concurrently within mating cell populations, and rDNA amplification precedes endoreplication by 1-2 h. Unexpectedly, our data indicate that the rDNA minichromosome is only transiently amplified in starved mated cells. It is subsequently constrained to replicate at a comparable rate as non-rDNA chromosomes upon re-feeding. The switch from rDNA amplification to rDNA endoreplication might be sensitive to the abundance of ORC or a labile amplification-specific trans-acting factor, possibly a protein that is bound to a region that creates a virtually impermeable replication fork barrier whose strength diminishes over time (Zhang et al., 1995). Dynamic changes in rDNA replication coincide with the previously reported down-regulation of ORC and MCM proteins in the developing macronucleus (Lee et al., 2015).

Several studies in mammals and flies raise the possibility of an ORC-independent DNA replication program. Homozygous *D. melanogaster* Orc1^-/-^ salivary gland cells endoreplicate to comparable levels as their wild type counterparts (10 endocycles) (Park and Asano, 2008). Similar results were obtained for the Orc2 k43^γ4^ allele. Disruption of the ORC 1 gene in diploid mouse tissues blocked DNA replication. However, ORC1 is not required in polyploid extraembryonic trophoblasts and hepatocytes Okano-Uchida et al., 2018). Finally, p53^-/-^ HCC116 human colon cancer cells initiate DNA replication in the absence of ORC1, ORC2 or ORC5 subunits (Shibata et al., 2016; Shibata and Dutta, 2020). Whether the site for MCM2-7 recruitment is marked by an aberrant DNA structure (hairpin or quadraplexe) or an alternative nucleation factor (non-ORC protein, non-coding RNA, stable RNA-DNA hybrid) remains to be determined. Compelling arguments for ORC-independent DNA replication in HU-arrested and released vegetative Tetrahymena (Sandoval et al., 2015), are consistent with the relaxed requirements for ORC during macronuclear development.

Finally, in this study, we determined that aberrant RIs generated during rDNA endoreplication emanate from a relatively small region, upstream of the rRNA transcription start site (Figure 4). Inactivation of the known 5’ NTS origins has been observed in ORC mutants that are defective in rDNA origin binding (Mohammad et al., 2007) and in HU-treated Tetrahymena, where ORC protein levels are reduced 50-fold (Sandoval et al., 2015). In both cases, an amorphous, alternate site was used analogous to what was observed in this study of endocycling wild type Tetrahymena and the vegetative TXR1 mutant. Since the rRNA coding region is unaffected in TXR1 knockout and endoreplicating wild type strains (Figure 4), rRNA transcripts are not implicated. The most likely source for aberrant RI production is the type 2 elements array, comprised of thirteen near perfect tandem 21 bp repeats (type 2 a-m) of unknown function. Type 2 elements are located between Domains 1 and 2 (the conventional 5’NTS origins) and the rRNA promoter (Figure 4A). Transient DNA melting of this AT-rich region could lead to out of register Watson:Crick strand reassociations that would produce unpaired single strand gaps or hairpin structures on opposite DNA strands-analogous to structures that are proposed for regions of the human genome that encode trinucleotide repeats (reviewed in Polleys, House and Freudenreich, 2017). The prediction is that two similar structures would form on each molecule, one on each strand-somewhat like the bidirectional replication fork. Could these hypothetical structures serve as molecular beacons for the DNA replication machinery when ORC is rate-limiting?

For those interested in pursuing gene expression, preliminary raw and processed RNA-seq datafiles on endoreplication phases I and II cells were previously deposited to the NCBI Gene Expression Omnibus (GEO; https://www.ncbi.nlm.nih.gov/geo/) under accession number GSE210103, along with cell cycle analysis of the vegetative cell cycle (Zhang et al., 2022).

## AUTHORS CONTRIBUTIONS

Xiangzhou Meng: experimental design, execution and interpretation, manuscript writing and graphics. Hung Quang Dang: experimental design, execution and interpretation, manuscript editing and graphics. Geoffrey Kapler: experimental design and interpretation, manuscript writing, graphics and funding.

## DATA ACCESS

All relevant data can be found within this paper and accompanying supplementary figures.

## ACKNOWLEDGEMENTS

We thank Doug Chalker for providing the histone H3-gfp tagged strain. This work was supported by NSF grant MCB-0132675 and the Tom and Jean McMullin Endowed Professorship to GMK.

---

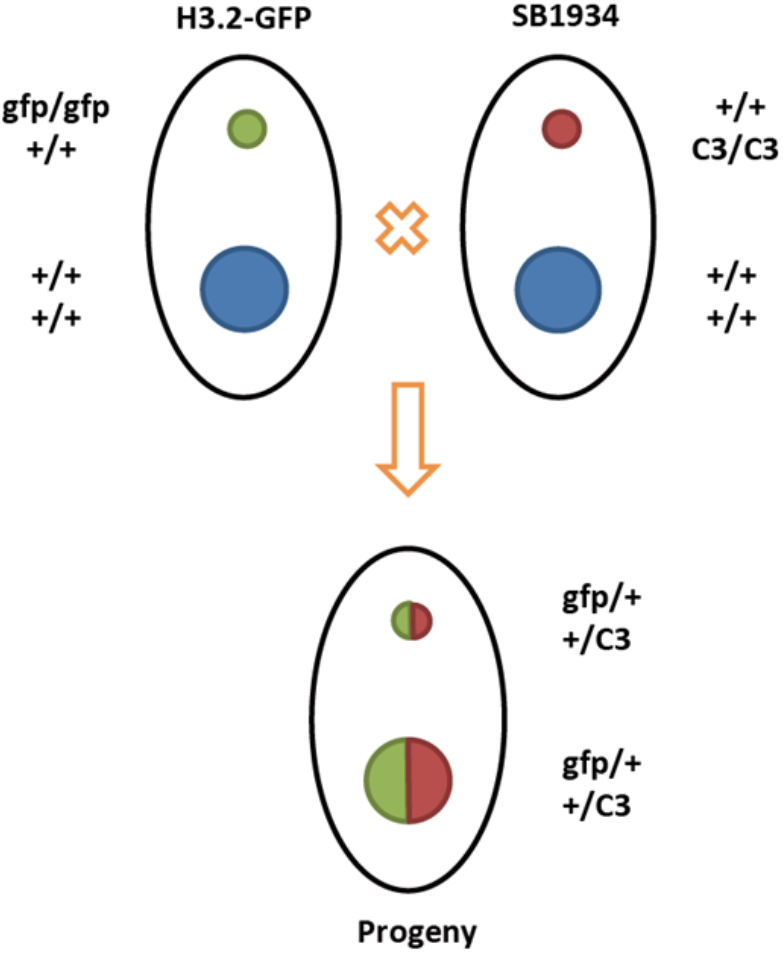

**Supplemental Figure 1**. Flow cytometry profile at 1h intervals during Endoreplication Phase 1 (cross: SB1943 x SB 4204).

**Supplemental Figure 2.** Schematic of mating between the two heterokaryons strains SB1934 (homozygous C3 rDNA micronucleus, B rDNA macronucleus) and H3.2-GFP (homozygous H3.2-GFP tagged micronucleus, wild type H3.2 macronucleus).

